# Incidental eagle carcass detection can contribute to fatality estimation at operating wind facilities

**DOI:** 10.1101/2022.10.21.513288

**Authors:** Eric Hallingstad, Daniel Riser-Espinoza, Samantha Brown

## Abstract

Risk of birds colliding with wind turbines, especially protected species like bald eagle and golden eagle, is a fundamental wildlife challenge the wind industry faces when developing and operating projects. The U.S. Fish and Wildlife Service requires wind facilities that obtain eagle take permits to document permit compliance through eagle fatality monitoring. If trained Operations and Maintenance (O&M) staff can reliably detect and report carcasses during their normal routines, then their ‘incidental detections’ could contribute substantially to meeting monitoring requirements required by eagle take permits. The primary objective of this study was to quantify incidental detection of eagle carcasses by O&M staff under a variety of landscape contexts and environmental conditions throughout 1 year. We used the incidental detection probabilities (proportion of decoys detected by O&M staff), along with raptor carcass persistence data and area adjustments, to calculate overall probability of incidental detection (i.e., incidental *g*). We used feathered turkey decoys as eagle-carcass surrogates for monthly detection trials at 6 study sites throughout the U.S. We evaluated the primary drivers of incidental detection using logit regression models including season, viewshed complexity, and a derived variable called the “density quartile” as covariates. We used an Evidence of Absence-based approach to estimate the overall probability of incidental detection. Detection probabilities decreased as viewshed complexity increased and as distance from the turbine increased. The resulting overall probability of incidental detection for the 12-month period ranged from 0.07 to 0.47 (mean = 0.31). The primary drivers of variability in incidental *g* were detection probability and the area adjustment. Results of our research show that O&M staff were capable of incidentally detecting trial carcasses while performing their typical duties. Incorporating incidental detection by O&M staff in eagle fatality monitoring efforts is a reliable means of improving estimates of a facility’s direct impacts on eagles.

## Introduction

Risk of birds colliding with wind turbines, especially sensitive or protected species, is a fundamental wildlife challenge the wind industry faces when developing and operating facilities [1]. Bald eagles (*Haliaeetus leucocephalus*) and golden eagles (*Aquila chrysaetos*) are susceptible to collision with wind turbines [2-4], and are protected under federal law [5]. Wind energy companies that obtain eagle take permits (ETPs), per the revised eagle permitting rules [6], are required by the U.S. Fish and Wildlife Service (USFWS) to complete eagle fatality monitoring to document permit compliance. Some level of eagle fatality monitoring will remain a requirement for all ETPs issued under the recently proposed revisions to the eagle permitting rules [7]. Given a permit term up to 30-years and a need to assess take compliance in 5-year increments, monitoring will be long-term with potentially high cost implications.

Eagle fatality monitoring has traditionally been completed by third parties hired to conduct standardized transect searches; however, over half of all reported eagle fatalities at wind energy facilities from 1997 to 2012 were detected incidentally outside of these standardized searches (i.e., the fatalities were detected by a property owner or by facility employees during routine site operations [3]). Incidental detections likely occur because eagle carcasses are large, tend to be highly visible, have long persistence times [8-11], and because Operations and Maintenance (O&M) staff, while traveling access roads and performing their typical duties on a near daily basis, have a more consistent presence on site than a third party. Furthermore, because O&M staff are visiting all turbines on a regular basis, there is likely more comprehensive spatial coverage of a facility compared to third-party monitoring that has traditionally been limited to a subset of turbines, and maximizing spatial coverage is desirable when monitoring for rare events such as eagle fatalities [12]. If O&M staff are educated on the importance of being aware of and reporting of potential eagle fatalities, and receive training to reliably detect and report carcasses found during their regular work schedule, eagle carcasses detected incidentally by O&M staff at some wind facilities could provide additional valuable information for compliance monitoring required by ETPs.

Accurate fatality estimates based on monitoring (whether standardized or incidental) require an adjustment for the overall probability of detection [13,14]. Although a number of fatality estimation models exist, Evidence of Absence (EoA; [15]) is a tool often used to estimate eagle fatality rates to determine permit compliance. EoA is specifically designed for fatality estimation in the context of “rare events” (i.e., 0 or very few carcasses are expected to be found during each search), which is the expectation for eagle fatalities at most wind energy facilities. EoA requires that an estimate of the overall probability of detecting an eagle carcass (i.e., *g*) is calculated for the period of interest, which incorporates estimate uncertainty and results in an estimate of eagle fatalities that may be higher than the number of observed eagle carcasses. Lower *g* results in higher uncertainty in the fatality estimate; conversely, a higher *g* will result in a more precise fatality estimate that is closer (or equal to) the observed count of carcasses (including 0).

Typical standardized fatality monitoring studies implemented by a third party have used EoA to estimate a *g* and a fatality estimate for eagles; however, the contribution of incidental detection has been overlooked to date, and an incidental *g* has not yet been quantified for wind facilities. In several ETPs recently issued by the USFWS, a *g* has been required in all years covered under an ETP [16-19]. Quantifying incidental detection and including its contribution in fatality monitoring plans will allow permittees to increase *g* over the permit term, develop a more accurate estimate of impacts to eagles, and provide the most robust means to demonstrate compliance.

Using incidental detection in eagle take estimation requires the information necessary to calculate *g*: bias correction factors include carcass *detection probability* (what is typically referred to as “searcher efficiency” in post-construction fatality monitoring), *carcass persistence* (the probability of carcasses persisting between detection opportunities), and an *area adjustment* (the proportion of carcasses present within the searched areas, and the proportion of turbines where detection was possible [i.e., sampled turbines]). The primary objective of this study was to quantify incidental detection of an eagle carcass surrogate by O&M staff during their normal maintenance activities under a variety of landscape contexts and environmental conditions throughout 1 year. Our secondary objective was to use the incidental detection probabilities from each site, along with area adjustments and relevant raptor carcass persistence data, to calculate incidental *g*’s that could contribute to an improved understanding of a facility’s direct impacts on eagles.

## Study areas

Five different USFWS Regions were selected (Regions 1, 2, 5, 6, and 8; Fig 1 [20-22]) for field trials to help capture a range of typical wind facility characteristics across the U.S. (Fig 2) and determine the variability of incidental eagle carcass detection across wind energy facilities. Geographic location, facility size (number of turbines), ground conditions, topography, and O&M staff activity on site (i.e., turbine visitation schedules) were all considered when evaluating candidate wind facilities for inclusion in this study. Field trials were conducted at 6 different wind energy facilities (study site[s]). General characteristics of each study site are provided in Table 1.

**Table 1.** Study site descriptions for the incidental eagle carcass detection study^a^.

| **Study Site** | **USFWS Region** | **Megawatts** | **Number of Turbines** | **Commercial Operation Date** | **USEPA Ecoregion (Level IV)** | **Primary Land Use(s)** | **Primary Land Cover(s)** | **Turbine Visitation Schedule** |
| --- | --- | --- | --- | --- | --- | --- | --- | --- |
| **Frontier I** | 2 | 200 | 61 | December 2016 | Prairie Tableland [23] | Cropland | Cropland | Monthly inspections |
| **Marble River** | 5 | 215 | 70 | October 2012 | Ontario Lowlands [24] | Cropland and livestock grazing | Forest and agriculture | Monthly inspections, unless inaccessible due to snow cover |
| **Mountain Wind I and II** | 6 | 141 | 67 | September 2010 | Rolling Sagebrush Steppe [25] | Livestock grazing | Sagebrush-steppe and grassland | Monthly inspections, unless inaccessible due to snow cover |
| **Pinyon Pines I and II** | 8 | 300 | 100 | December 2012 | Western Mojave Basin [26] | Livestock grazing | Desert shrub | Routine inspections varied due to weather conditions; approximately monthly |
| **Shiloh I** | 8 | 150 | 100 | April 2006 | Suisun Terraces and Low Hills [27] | Livestock grazing and cropland | Grassland and cropland | Monthly inspections |
| **Wild Horse** | 1 | 273 | 127 | December<br>2006 | Yakima Folds<br>and Okanogan<br>Drift Hills [28] | Livestock<br>grazing | Shrub-steppe<br>and grassland | Monthly inspections |

<sup>a</sup> Abbreviations: USFWS = U.S. Fish and Wildlife Service ,USEPA = U.S. Environmental Protection Agency

**Fig 1.**
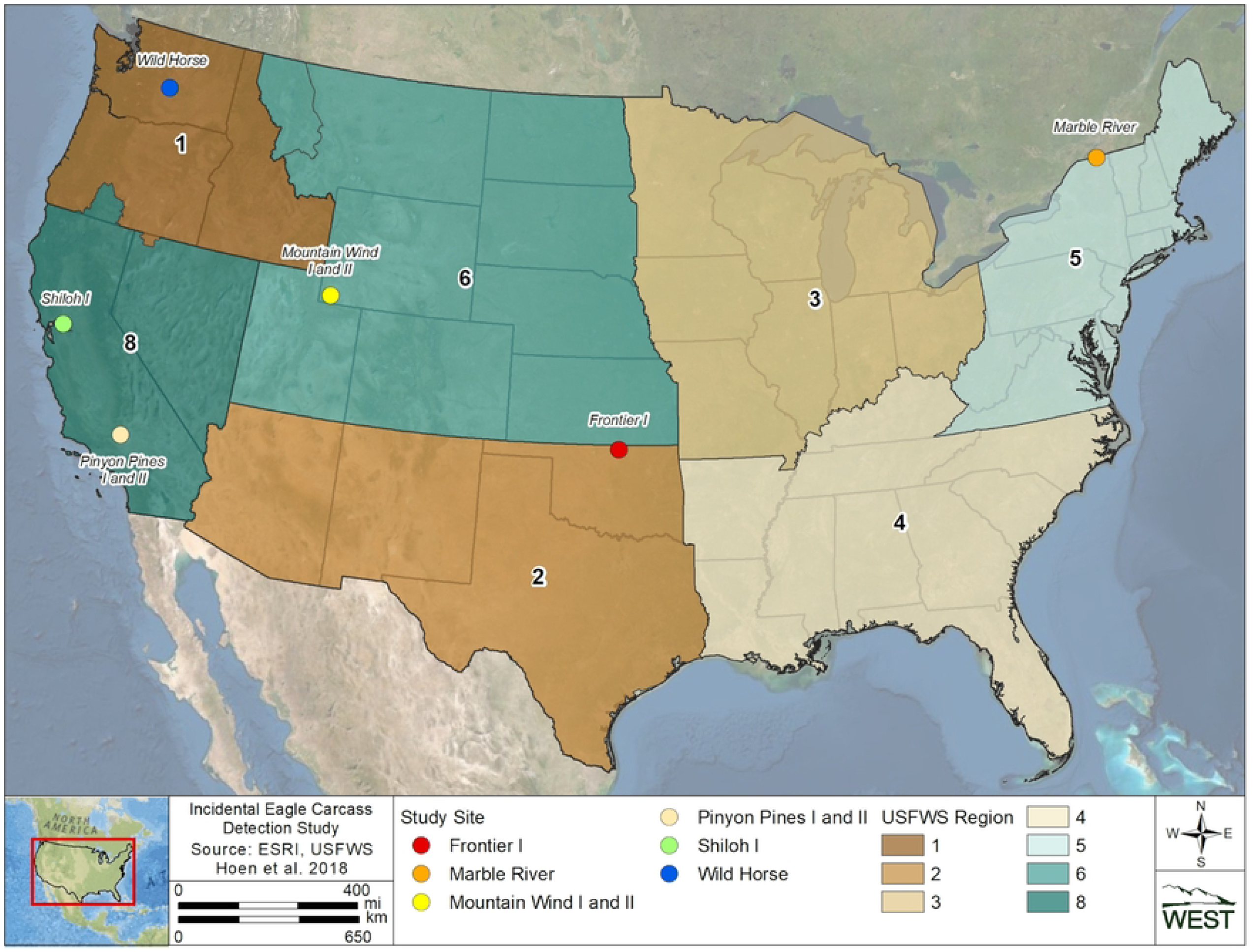
Study site locations included in the incidental eagle carcass detection study conducted from June 27, 2021, through July 14, 2022.

**Fig 2.**
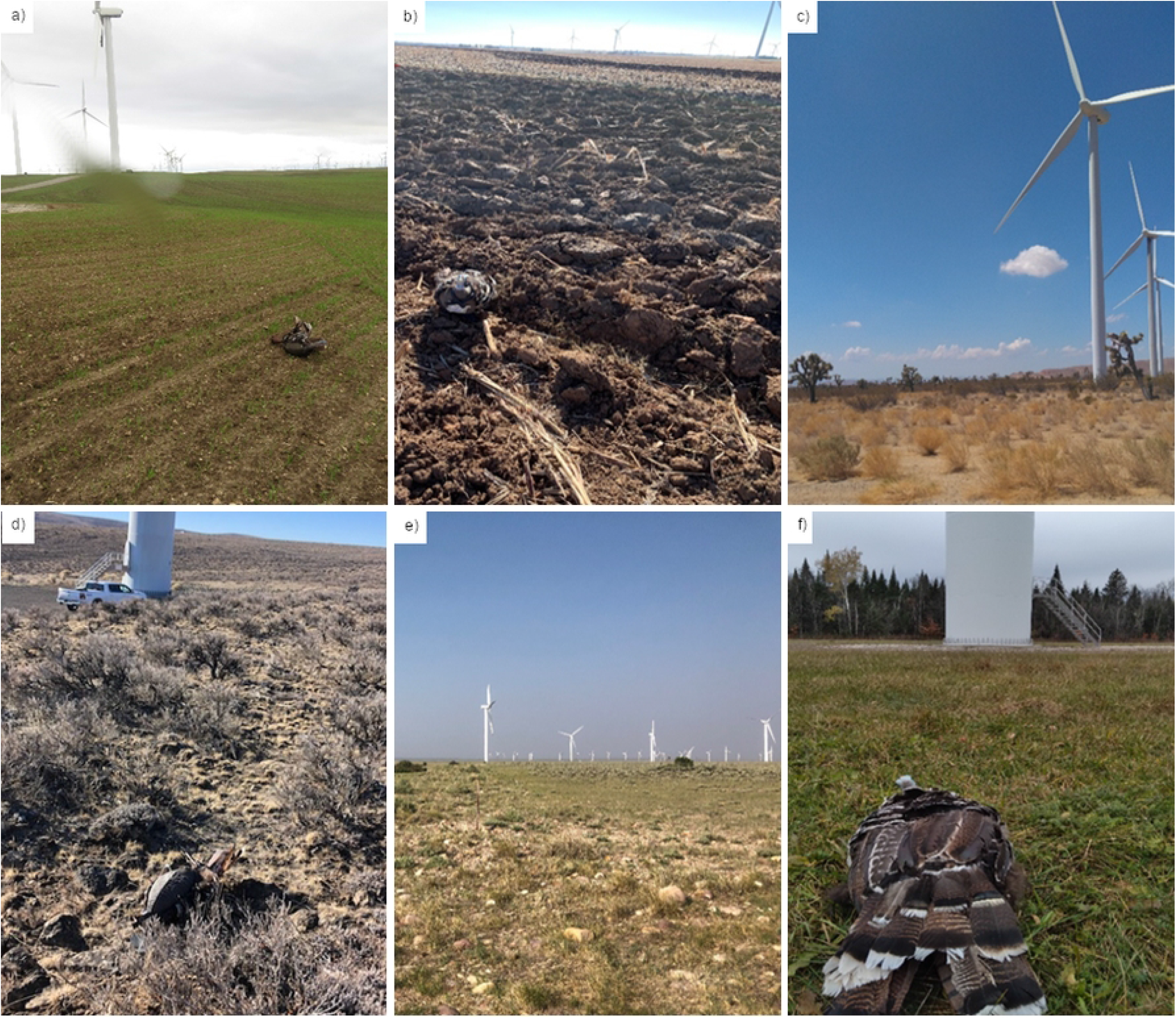
Examples of variable land cover types present at the wind energy facilities included in this study. The study sites shown in the photographs are (a) Shiloh I, (b) Frontier I, (c) Pinyon Pines I and II, (d) Wild Horse, (e) Mountain Wind I and II, and (f) Marble River. Decoy placements are visible in several photographs.

## Methods

### Viewshed complexity mapping

We expected detection probability to be influenced by viewshed complexity (i.e., the height/density of vegetation and variation in topography) that could conceal carcasses from view within the searchable area at each turbine. Therefore, prior to the field trials, we mapped the viewshed within the search area, defined as a 100-m radius circle centered on every turbine, at all 6 study sites. We categorized viewshed complexity as low, moderate, or high complexity based on ground cover conditions shown in Table 2. We also characterized areas as unviewable/unsearchable when visibility out to 100 m was limited by variation in terrain, mature crops, or forest. Areas categorized as unviewable/unsearchable were not visible by O&M staff conducting regular maintenance duties, such as when traveling in a vehicle to a turbine or when they were standing near turbine bases or edges of turbine pads. Based on the viewshed mapping performed at the beginning of our study and the ground cover conditions present within the search areas, we inferred predominant viewshed complexity classes by season at each study site. For example, we assumed cropland areas typically progressed from low to moderate to unviewable/unsearchable across the crop-growing cycle.

**Table 2.** Viewshed complexity classifications. Table 2 legend. Classifications were based on percent bare ground and vegetation height within the 100-m radius search areas centered on wind turbine towers during detection trials conducted at the study sites from June 27, 2021, through July 14, 2022.

| Percent Bare Ground | Average Vegetation Height in Non-Bare Ground Areas |  |  |  |  |
| --- | --- | --- | --- | --- | --- |
|  | <15 cm | 16-30 cm | 31-45 cm | > 46 cm | Rocky |
| > 90% | Low | Low | Low | Low | Moderate |
| 26-89% | Low | Moderate | Moderate | Moderate | Moderate |
| 1-25% | Low | Moderate | High | High | High |
| 0% | Low | Moderate | High | High | High |

We also classified viewshed complexity for each decoy placed during our detection trials. Classifications were made based on the same parameters described above, but limited to ground cover within a 5-m radius around each placement. By determining real-time viewshed complexities associated with each opportunity for detection, we were able to evaluate the effects of viewshed complexity on detection probabilities.

### Field methods for incidental detection trials

We used Turkey Skinz™ (A-Way Hunting Products, Beaverton, Michigan) feathered turkey (*Meleagris gallopavo*) decoys as eagle carcass surrogates for all detection trials. Decoys were used to approximate the size and color (and, therefore, detectability) of an eagle carcass, and were also a practical choice to obtain adequate sample size and ensure trials were still in place when the opportunity for detection occurred. Decoy placements were assigned a random distance and bearing from the base of randomly selected turbines throughout the study period. Decoy placement distances were randomized from a distribution of raptor fall distances derived from a meta-analysis [10], truncated to 100 m from the turbine base, which captures approximately 94.5% of expected raptor fatalities for the turbine sizes at our study sites. Thus, the density of decoy placements increased from 0 to 40 m from the turbine base and then decreased out to 100 m [10]. Placement bearing-from-turbine was chosen from a uniform distribution between 0 and 360 degrees. To the extent possible, placements were stratified across the mapped viewshed complexity categories (low, moderate, high). No decoy was placed in an unviewable/unsearchable area.

We deployed up to 16 decoys per study site per month, for approximately 48 decoys at each study site per quarter. A larger monthly sample size was initially planned, but we reduced our sampling effort due to concerns expressed by USFWS regarding high decoy densities biasing O&M staff detection probabilities. The rate at which decoys were placed at sites was not intended to accurately mimic a realistic eagle fatality rate because our study was constrained by time, cost, and sample size requirements. Nonetheless, even at our smallest study sites, decoys were only placed at roughly 25% of a facility’s turbines during each monthly trial and not all decoys were detected (see Results section, below); thus, O&M staff were not detecting nor were decoys present at the majority of turbines O&M staff visited in any given month. All decoys were placed on a single day each month unless site conditions or other field delays required placements spanning 2 days. Western EcoSystems Technology, Inc. (WEST), personnel placed the decoys without the direct knowledge of the O&M staff, and most placements occurred on weekends or after hours when O&M staff were not present.

Decoys were left in place for 1 month or until they were detected, whichever was sooner. One month was chosen as our trial length, as this timeframe aligned with a common visitation schedule for O&M staff making periodic visits to all study site turbines. As such, we assumed each decoy had at least 1 opportunity for detection during each round of placements. Prior to field trials, O&M staff were briefed on the objectives of the study and instructed on detection and documentation protocols, but were directed to not otherwise deviate from their typical maintenance routines. When a decoy was detected by O&M staff, the decoy was immediately collected from the field, deposited at the O&M building, and reported by O&M staff using a simple decoy detection form. O&M staff were asked to record the date, nearest turbine, their activity when the detection was made, and the decoy tag identification.

Seasonal adjustments to deployment schedules were needed at several study sites. During the winter months, roads were not cleared of snow at Marble River, Mountain Wind I and II, and Wild Horse, and on-site travel was limited to as-needed maintenance visits. Furthermore, frequent snowfall was anticipated to obscure decoy presence on the landscape at those 3 study sites. Opportunities for decoy detection were expected to be minimal under these conditions, so no decoys were placed from December 2021 – March 2022 at Marble River, from January – March 2022 at Mountain Wind I and II, and from December 2021 – January 2022 at Wild Horse. In 2 cases, winter weather conditions also resulted in abbreviated trials. At Mountain Wind I and II, decoys placed on November 30, 2021, were retrieved on December 14, 2021, ahead of a predicted snowstorm, approximately 2 weeks short of the standard trial length. At Wild Horse, decoys placed on November 30, 2021, became completely covered in snow after an unexpected snowstorm on December 22, 2022. Despite the shortened trial length, the number of decoys detected by O&M staff before being covered by snow was consistent with the number of decoys detected in previous months at these sites. Thus, we assumed the decoys were placed ahead of all or most O&M visits to turbines that month and these placements should be included in analysis despite the abbreviated trial period potentially limiting additional opportunities for detection.

We also anticipated decoy detection would be negligible in cropland areas when crop heights exceeded the size of the carcass. Therefore, no deployments were made in cropland areas during the summer growing season. All areas temporarily or permanently excluded from the monthly trial schedule were accounted for in the area adjustment component when calculating the overall probabilities of detection (see Area adjustment section, below). Specifically, 24 turbines at Frontier I and 41 turbines at Shiloh I were treated as unsearchable cropland during the summer season. We considered all major changes in land cover due to snowfall or agricultural activity (e.g., harvest time) when determining season dates for each study site (Table 3), as these changes were expected to impact detection probability. We attempted to make seasons equal in length to the greatest extent practicable.

**Table 3.** Season dates for the incidental detection study.

at the 6 study sites from June 27, 2021, to July 14, 2022.
| Site | Season Dates |  |  |  |
| --- | --- | --- | --- | --- |
|  | Spring | Summer | Fall | Winter |
| **Frontier I** | March 16 –<br>June 15 | June 16 – August<br>31 | September 1 –<br>December 15 | December 16 –<br>March 15 |
| **Marble River** | April 1 – June<br>25 | June 26 – August<br>30 | August 31 –<br>November 30 | December 1 –<br>March 31 |
| **Mountain Wind**<br>**I and II** | April 1 – June<br>30 | July 1 –<br>September 20 | September 21 –<br>November 30 | December 1 –<br>March 31 |
| **Pinyon Pines I**<br>**and II** | March 1 – May<br>31 | June 1 – August<br>31 | September 1 –<br>November 30 | December 1 –<br>February 28 |
| **Shiloh I** | April 1 – June<br>30 | July 1 –<br>September 30 | October 1 –<br>December 31 | January 1 – March<br>31 |
| **Wild Horse** | March 1 – May<br>31 | June 1 – August<br>31 | September 1 –<br>November 30 | December 1 –<br>February 28 |

Other conditions at some study sites made decoy placements at some turbines infeasible or inappropriate for the objectives of our study. For example, Marble River and Frontier I had landowners who did not permit decoy placements on their land during the entire study. This restriction excluded 12 turbines at Marble River and 1 turbine at Frontier I. An additional 6 turbines at Shiloh I did not receive placements during 1 month in spring because the property was not accessible. Finally, 10 turbines at Shiloh I were subject to dedicated wildlife monitoring in the spring and fall, so no decoys were placed at these turbines to avoid biasing the results with detections from standardized searches. Turbines at which decoys could not be placed were otherwise assumed to be similar to other turbines at the same study site with respect to viewshed complexity and the visitation schedule by O&M staff.

### Analysis methods

#### Incidental detection trials

We summarized incidental decoy detection data by study site, viewshed complexity class, and season using only decoys determined to be available for detection. Decoys were considered available unless they were not found by O&M staff and could not be recovered by the trial administrator following the month-long trial period. Decoys detected by O&M staff during the month-long trial period after placement were considered “found”; any decoys collected after the month-long trial period were treated as “available and not found.” We also summarized the available self-reported detection data from O&M staff to determine general patterns in the activities being performed when detections occurred. Since O&M data were not reported consistently at each site throughout the entire year, we did not attempt to incorporate these data into any further modeling efforts.

We used the incidental detection trial data in 2 ways to accomplish different goals. The first use of incidental detection trial data was exploring spatial and temporal patterns across the 6 study sites, with the goal of providing inference about the drivers of detection. The second use of the incidental detection trial data was to calculate site-specific incidental *g*’s in the manner expected for a single site evaluating compliance with permit conditions. The second application of incidental detection trials is described in more detail below (see Calculating the overall probability of incidental detection [incidental *g*]).

To explore potential drivers (i.e., covariates) of incidental detection, trial data were pooled across study sites and used to fit logit regression models, as implemented in Dalthorp et al. [14]. Covariates included in model selection were season, viewshed complexity, and a derived variable called the “density quartile”. The density quartile provides a simplified way to associate decoys with their proximity to a turbine by creating distance bins that align with the expected density of raptor carcasses on the landscape (near: 0–33 m; near-mid: 34–45 m; far-mid: 46–61 m; and far: more than 61 m from turbine base; using carcass distribution in Hallingstad et al. [14]). For example, the first density quartile “near” is where we would expect the first 25% of large raptor fatalities to fall relative to a turbine.

In calculating incidental *g*, we also considered the possibility that a real eagle carcass that goes undetected during 1 search might remain available for detection on subsequent searches. A carcass missed at least once likely has a lower detectability than a carcass that was not missed, possibly because the carcass was in a less conspicuous location, because it was decaying between searches, or for other reasons. This detection reduction factor (*k*) can range from 0 to 1, where *k* equal to 0 implies a carcass missed on the first search opportunity would never be found on subsequent searches. A *k* of 1 implies detection probability remained constant no matter how many times a carcass was missed. Estimating *k* is difficult because it requires the placement and tracking of a large number of trial carcasses through multiple searches. We did not estimate *k* for our analysis, but rather assumed a value of 0.67 since it is currently the only published estimate of *k* and consistent with a previous study of fatality monitoring for eagles and large raptors [14,29].

We fit logit regression models using all possible combinations of the 3 potential covariates and 2-way interactions, and fixed *k*. We dropped models with interactions where at least 1 of the interaction covariates was not included as a main effect. Model selection was performed using an information theoretic approach known as AICc, or corrected Akaike Information Criterion [30]. The best model was selected as the most parsimonious model within 2 AICc units of the model with the lowest AICc value.

#### Carcass persistence

Estimating the average probability of persistence is necessary to calculate *g* using EoA. When estimating eagle or other large raptor (i.e., raptors with a minimum 30-cm wing chord and 300-g mass) fatality rates, raptor persistence data should be used whenever possible; game birds (e.g., mallards [*Anas platyrhynchos*], ring-necked pheasants [*Phasianus colchicus*]) consistently have shorter persistence times than large raptor carcasses (e.g., red-tailed hawks [*Buteo jamaicensis*], great horned owls [*Bubo virginianus*]), which would result in a lower probability of persistence and an inflated fatality estimate [11]. However, site-specific raptor persistence data were not available for 4 of our 6 study sites. For Wild Horse, we used existing raptor persistence data collected on 58 large raptor carcasses placed between May 2016 and September 2020. For Shiloh I, we used 35 large raptor carcasses placed at adjacent facilities in the Montezuma Hills Wind Resource Area (S1 Table) between March 2012 and October 2013. For the remaining 4 study sites, we used large raptor persistence data from several other wind energy facilities that were comparable in location and/or site characteristics [11].

We fit interval-censored survival regression models [11,14, 31-33] to each dataset to generate the parameter estimates required by EoA to account for persistence probability in *g*. We compared several candidate models for each study site, including different underlying persistence distributions and potential covariates. Candidate models were fit using exponential, log-logistic, lognormal, and Weibull survival distributions to characterize a broad range of persistence dynamics. Season was the only potential covariate considered in models, with the exception of Shiloh I, where there was not a sufficient sample size across seasons; thus, intercept-only models were considered for Shiloh I. We fit models that included potential covariates (where relevant) on the location and scale parameters used to define the distributions above; see Kalbfleisch and Prentice [32], Dalthorp et al. [14,21] and Therneau [33] for details about the location and scale parameterizations used for the candidate survival distributions. Model selection was performed using an information theoretic approach known as AICc [30]. The best model was selected as the most parsimonious model within 2 AICc units of the model with the lowest AICc value. The parameter estimates of the selected model (α [shape] and β [scale], including the 95% confidence interval [CI] of β) were used as inputs in the EoA Single Class Module. See Hallingstad et al. [11] for additional details about the raptor persistence data.

#### Area adjustment

The area adjustment component of *g* accounts for the amount of area within the nominal search region (in this case, 100 m) and the expected occurrence, or density, of carcasses on the landscape. The density-weighted area adjustment was estimated as the product of the viewable area around each turbine and a carcass-density distribution. The carcass-density distribution predicts the likelihood a carcass falls a given distance from the turbine base. At the study sites with restricted search areas (due to snowfall in winter and croplands at peak growing season), the amount of viewable area and/or proportion of searchable turbines were reduced based on the length of time carcass detection was not feasible.

The raptor carcass-density distribution from Hallingstad et al. [10] was used to calculate the searched area adjustment in this study. The density distribution developed in Hallingstad et al. [10] is based on a meta-analysis of raptor spatial data from multiple wind facilities with varying turbine designs and wind regimes.

#### Calculating the overall probability of incidental detection (incidental *g*)

Estimating *g* depends on the carcass detection probability (i.e., the proportion of available decoys/carcasses found, and the factor by which detection probability decreases on subsequent searches [*k*]), carcass persistence (the probability of carcasses persisting between detection opportunities), and an area adjustment (the proportion of carcasses present within the searched areas, and the proportion of turbines where detection was possible [i.e., searched turbines]). We used the EoA [15] modeling framework to estimate the incidental *g* at each study site resulting from detection of decoys by O&M staff during the study period, the raptor persistence data most appropriate for each site, and the area adjustment information. The estimated overall probability of a carcass being both available and incidentally detected (i.e. incidental *g*) can be approximated as

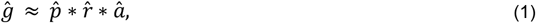

Where 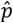 is average detection probability, 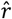 is the average probability of carcass persistence between detection opportunities, and â is the area adjustment (“hat” accents are added to notate estimated values). EoA allows *g* to be calculated for any strata or “class” (e.g., season within a site) using the Single Class Module, and then to combine *g* across all strata via the Multiple Class Module [15]. For each site, incidental detection trial effort and results could vary by season (based on changes in viewshed complexity over the course of a year), raptor persistence models could have seasonal covariates, and the area adjustment could vary based on changes in viewable area (e.g., due to crop cycles).Thus, we used the Single Class Module to calculate incidental *g* by season, then combined across seasons using the Multiple Class Module to arrive at a site-specific incidental *g*. We did not additionally stratify by viewshed complexity class (or density quartile) because incidental detection trials were placed at random distances from turbines based on the expected fall distribution of raptors [10] at each study site, and therefore the results of those trials would be representative of the viewshed complexity distribution unique to each study site. Consistent with our effort to model drivers of incidental detection probability, we assumed a *k* of 0.67 to estimate incidental *g*. Finally, based on known turbine maintenance visitation schedules, search effort was assumed to be 30 days (although many turbines would be visited [i.e., “searched”] more frequently during an average 30-day period) for the purposes of estimating persistence probabilities (i.e., the 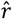 component of incidental *g*).

In calculating incidental *g*, we also considered a temporal component to eagle risk. Eagle use of some areas can vary throughout the annual cycle, limiting the risk of collision (and, therefore, the potential for an eagle fatality to occur) to certain seasons. For simplicity, temporal risk was assumed to be uniform across the study period as evaluating collision risk periods was beyond the scope of our study. All estimates were calculated using the EoA R package (version 2.0.7), using the Single Class Module and Multiple Class Module of EoA [15].

## Results

### Viewshed complexity mapping

Viewshed complexity classes were mapped in the field and digitized in ArcGIS [34]. The total area within 100 m of each turbine was aggregated by viewshed complexity class to determine the predominant viewshed complexity class within the study sites (Table 4). For cropland areas, the percentage of high-complexity search area fluctuated as cropland visibility varied on a seasonal basis. For example, cropland viewshed complexity at Frontier I was predominately low during the fall and winter, but cycled through moderate, high, and unviewable/unsearchable in the spring and summer. To a lesser extent, viewshed classification cycling occurred over the course of the growing seasons at Marble River and Shiloh I. Viewshed complexity within other land cover types remained relatively consistent from season to season. Reduced on-site travel during the winter season due to large amounts of snowfall is another mechanism through which viewshed complexity varied seasonally (Table 4).

**Table 4.** Predominant viewshed complexity classes.

legend. Complexity classes by season during detection trials conducted at the study sites from June 27, 2021, through July 14, 2022.
| Study Site | Predominant Viewshed Complexity Class |  |  |  |
| --- | --- | --- | --- | --- |
|  | Spring | Summer | Fall | Winter |
| **Frontier I** | Moderate | Unviewable/ unsearchable | Low | Low |
| **Marble River** | Moderate | Unviewable/ unsearchable | Moderate | Unviewable/ unsearchable |
| **Mountain Wind I and II** | Moderate | Moderate | Moderate | Unviewable/ unsearchable |
| **Pinyon Pines I and II** | High | High | High | High |
| **Shiloh I** | Moderate | Moderate | Low | Moderate |
| **Wild Horse** | Moderate | Moderate | Moderate | Moderate |

### Incidental detection trials by study site

Over the 12 month study, we placed 996 decoys in the field. Of the 996 decoy placements, 918 were confirmed as available for detection (either detected or collected at the end of the 30-day trial) and 444 of available decoys were detected by O&M staff (48%; Table 5); 78 decoy placements were unavailable for detection due to theft, agricultural activities, or undetermined means of removal. The probability of O&M staff detecting available decoys ranged from 0.28 (Wild Horse) to 0.78 (Marble River; Table 5).

**Table 5.** Incidental detection trial results by study site.

legend. Trial results for field studies during detection trials conducted at the study sites from June 27, 2021 through July 14, 2022.
| **Study Site** | **Decoys Placed** | **Decoys Available** | **Decoys Found** | **Average Detection**<br>**Probability** | **Monthly**<br>**Trials** |
| --- | --- | --- | --- | --- | --- |
| **Frontier I** | 177 | 140 | 63 | 0.45 | 12 |
| **Marble River** | 116 | 86 | 67 | 0.78 | 7 |
| **Mountain**<br>**Wind I and II** | 144 | 144 | 91 | 0.63 | 8 |
| **Pinyon Pines I**<br>**and II** | 192 | 188 | 59 | 0.31 | 12 |
| **Shiloh I** | 191 | 186 | 115 | 0.62 | 12 |
| **Wild Horse** | 176 | 174 | 49 | 0.28 | 11 |
| **Overall** | **996** | **918** | **444** | **0.48** | **57** |

Variation in incidental detection at the study sites was driven primarily by variation in viewshed complexity. Decoy detection in high viewshed complexity ranged from no detection (0 out of 8 placements detected) at Marble River and Wild Horse (0 out of 6 placements detected) to 0.31 at Frontier I (8 out of 26 placements detected; Table 6). In contrast, detection in low viewshed complexity ranged from 0.31 at Wild Horse (39 out of 126 placements detected) to 0.89 at Marble River (50 out of 56 placements detected), while detection in moderate viewshed complexity ranged from 0.10 at Frontier I (2 out of 22 placements detected) to 0.77 at Marble River (17 out of 22 placements detected; Table 6).

**Table 6.**
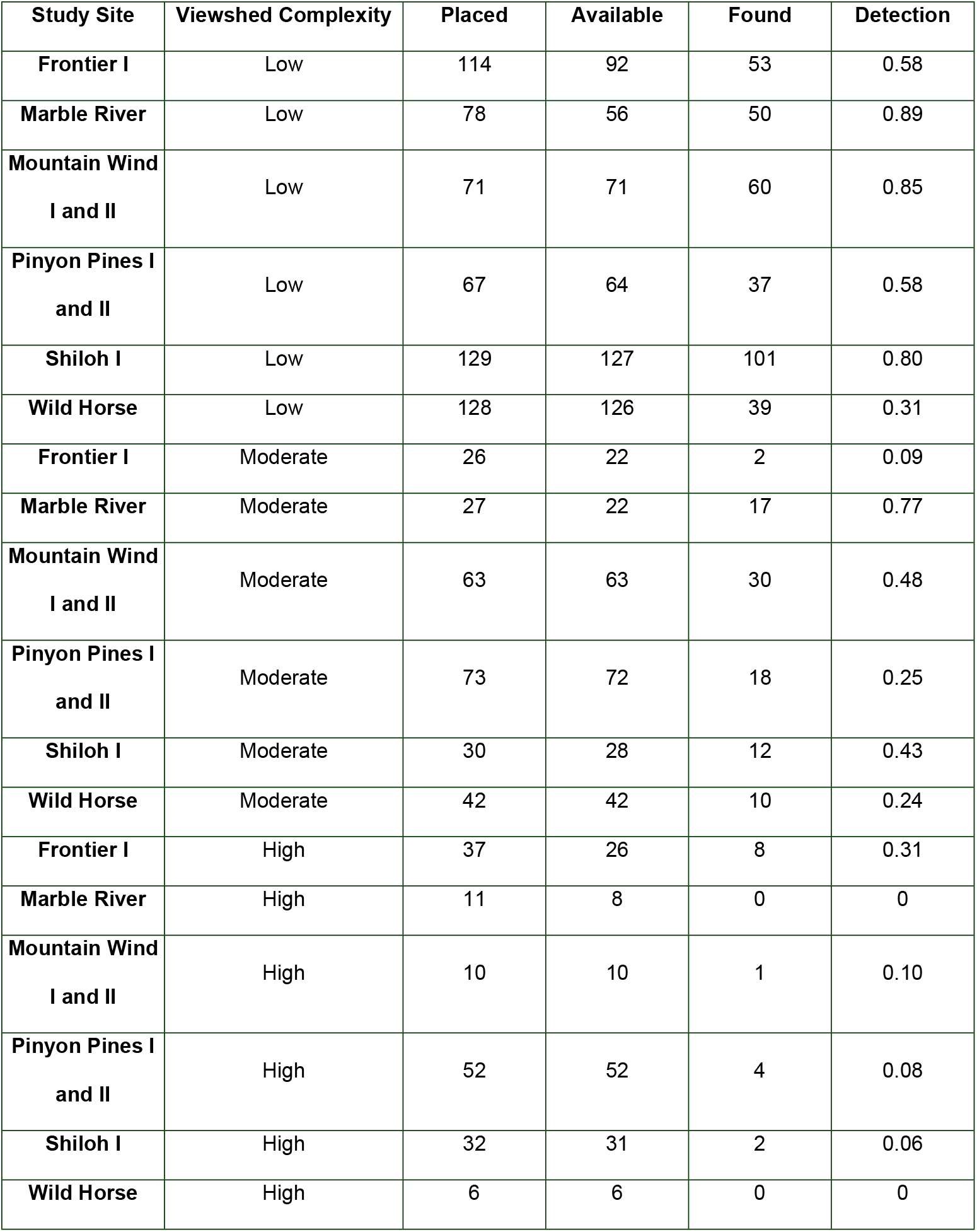
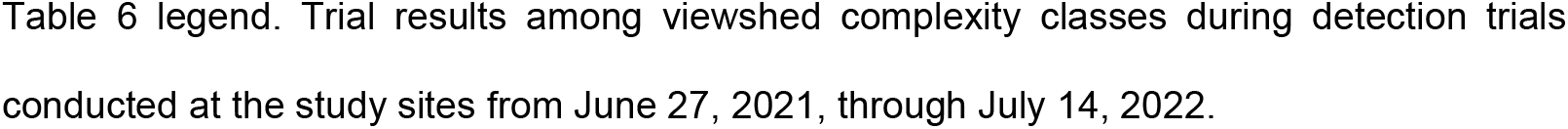
Incidental detection trial results among viewshed complexity classes.

legend. Trial results among viewshed complexity classes during detection trials conducted at the study sites from June 27, 2021, through July 14, 2022.
| **Study Site** | **Viewshed Complexity** | **Placed** | **Available** | **Found** | **Detection** |
| --- | --- | --- | --- | --- | --- |
| **Frontier I** | Low | 114 | 92 | 53 | 0.58 |
| **Marble River** | Low | 78 | 56 | 50 | 0.89 |
| **Mountain Wind I and II** | Low | 71 | 71 | 60 | 0.85 |
| **Pinyon Pines I and II** | Low | 67 | 64 | 37 | 0.58 |
| **Shiloh I** | Low | 129 | 127 | 101 | 0.80 |
| **Wild Horse** | Low | 128 | 126 | 39 | 0.31 |
| **Frontier I** | Moderate | 26 | 22 | 2 | 0.09 |
| **Marble River** | Moderate | 27 | 22 | 17 | 0.77 |
| **Mountain Wind I and II** | Moderate | 63 | 63 | 30 | 0.48 |
| **Pinyon Pines I and II** | Moderate | 73 | 72 | 18 | 0.25 |
| **Shiloh I** | Moderate | 30 | 28 | 12 | 0.43 |
| **Wild Horse** | Moderate | 42 | 42 | 10 | 0.24 |
| **Frontier I** | High | 37 | 26 | 8 | 0.31 |
| **Marble River** | High | 11 | 8 | 0 | 0 |
| **Mountain Wind I and II** | High | 10 | 10 | 1 | 0.10 |
| **Pinyon Pines I and II** | High | 52 | 52 | 4 | 0.08 |
| **Shiloh I** | High | 32 | 31 | 2 | 0.06 |
| **Wild Horse** | High | 6 | 6 | 0 | 0 |

Detection probability (decoys found divided by decoys available) also varied by season across study sites, with the greatest range in fall (0.16–0.94) and smallest range in winter (0.21–0.51; Table 7). Detection probabilities at Frontier I, Shiloh I, and Mountain Wind I and II were lowest in spring (0.24, 0.36, and 0.56, respectively). Detection probability at Marble River was lowest in summer (0.57). Detection at Frontier I was highest in winter (0.51), whereas at Shiloh I detection was highest in summer (0.86), and Marble River had the highest detection in fall (0.94). At Pinyon Pines I and II, detection was similar in spring and fall (0.30 and 0.33, respectively), but notably higher in summer compared to winter (0.44 and 0.21, respectively; Table 7). Wild Horse had higher detection probabilities in winter and spring (0.32 and 0.40, respectively) than in summer and fall (0.24 and 0.16, respectively; Table 7).

**Table 7.** Incidental detection trial results.

legend. Incidental results by study site and season for field studies during detection trials conducted at the study sites from June 27, 2021, through July 14, 2022.
| Study Site | Season | Decoys Placed | Decoys Available | Decoys Found | Detection Probability |
| --- | --- | --- | --- | --- | --- |
| Frontier I | Spring | 40 | 34 | 8 | 0.24 |
| Marble River | Spring | 41 | 31 | 24 | 0.77 |
| Mountain Wind I<br>and II | Spring | 64 | 64 | 36 | 0.56 |
| Pinyon Pines I<br>and II | Spring | 48 | 47 | 14 | 0.30 |
| Shiloh I | Spring | 47 | 47 | 17 | 0.36 |
| Wild Horse | Spring | 48 | 48 | 19 | 0.40 |
| Frontier I | Summer | 27 | 15 | 6 | 0.40 |
| Marble River | Summer | 32 | 23 | 13 | 0.57 |
| **Mountain Wind I and II** | Summer | 32 | 32 | 24 | 0.75 |
| **Pinyon Pines I and II** | Summer | 36 | 36 | 16 | 0.44 |
| **Shiloh I** | Summer | 64 | 64 | 55 | 0.86 |
| **Wild Horse** | Summer | 64 | 63 | 15 | 0.24 |
| **Frontier I** | Fall | 64 | 52 | 29 | 0.56 |
| **Marble River** | Fall | 43 | 32 | 30 | 0.94 |
| **Mountain Wind I and II** | Fall | 48 | 48 | 31 | 0.65 |
| **Pinyon Pines I and II** | Fall | 60 | 58 | 19 | 0.33 |
| **Shiloh I** | Fall | 32 | 31 | 22 | 0.71 |
| **Wild Horse** | Fall | 32 | 32 | 5 | 0.16 |
| **Frontier I** | Winter | 46 | 39 | 20 | 0.51 |
| **Marble River<sup>a</sup>** | Winter | 0 | 0 | 0 | 0 |
| **Mountain Wind I and II<sup>a</sup>** | Winter | 0 | 0 | 0 | 0 |
| **Pinyon Pines I and II** | Winter | 48 | 47 | 10 | 0.21 |
| **Shiloh I** | Winter | 48 | 44 | 21 | 0.48 |
| **Wild Horse** | Winter | 32 | 31 | 10 | 0.32 |

<sup>a</sup>Winter trials were not conducted at Marble River or Mountain Wind I and II due to the amount of snowfall and limited site access.

The activity being performed when a decoy detection occurred was recorded for 307 out of the 444 detections (69%, Table 8). Of these, the majority of detections occurred while O&M staff were at turbines conducting routine inspections (106 detections; 35%), followed closely by detections made while driving (96 detections; 31%) and while performing turbine maintenance (91 detections; 30%; Table 8). A small number of detections were made while performing other activities on-site, such as land management (e.g., weed control, mowing) or surveying (14 detections; 5%; Table 8).

**Table 8.** Incidental detection trial results by Operations and Maintenance staff activity.

legend. Trial results by activity being performed by Operations and Maintenance staff at the time of detection during the detection trials conducted at the study sites from June 27, 2021, through July 14, 2022.
| Study Site | Maintenance | Driving | Land Management | Turbine Inspections | Surveying | Total Reported | Total Found |
| --- | --- | --- | --- | --- | --- | --- | --- |
| Frontier I | 12 | 0 | 0 | 0 | 0 | 12 | 63 |
| Marble River | 12 | 3 | 10 | 2 | 0 | 27 | 67 |
| Mountain Wind I and II | 14 | 41 | 1 | 32 | 0 | 88 | 91 |
| Pinyon Pines I and II | 27 | 13 | 0 | 1 | 0 | 41 | 59 |
| Shiloh I | 11 | 26 | 0 | 68 | 0 | 105 | 115 |
| Wild Horse | 15 | 13 | 0 | 3 | 3 | 34 | 49 |
| **Total** | **91** | **96** | **11** | **106** | **3** | **307** | **444** |

#### Modeling drivers of incidental detection

The best-supported model of incidental detection included main effects for density quartile and viewshed complexity (Delta [Δ] AICc 0.37; Table 9). The other 9 top-ranked models all included main effects for density quartile and viewshed complexity, suggesting these 2 covariates are relatively more important for explaining variability in incidental detection compared to season alone or season in conjunction with 1 of the other 2 covariates. Although incidental detection varied by season with study sites (Table 9), season did not rank as important as viewshed complexity and density quartile (which is a surrogate for distance from turbine) in explaining variation in incidental detection across the entire dataset.

**Table 9.** Model selection results for incidental detection.

legend. Results using AICc for 18 models of incidental detection using 918 decoy placements during detection trials conducted at 6 study sites from June 27, 2021, through July 14, 2022.
| **Covariates** | **Number of**<br>**Parameters** | **AICc<sup>a</sup>** | **$\Delta$ AICc<sup>b</sup>** |
| --- | --- | --- | --- |
| **Season + Viewshed Complexity + Density Quartile + Season: Viewshed Complexity** | 15 | 1097.68 | 0 |
| **Season + Viewshed Complexity + Density Quartile** | 9 | 1097.76 | 0.08 |
| **Viewshed Complexity + Density Quartile** | 6 | 1098.05 | 0.37<sup>c</sup> |
| **Viewshed Complexity + Density Quartile + Viewshed Complexity: Density Quartile** | 12 | 1101.43 | 3.75 |
| **Season + Viewshed Complexity + Density Quartile + Viewshed Complexity: Density**<br>**Quartile** | 15 | 1102.16 | 4.48 |
| **Season + Viewshed Complexity + Density Quartile + Season: Viewshed Complexity +**<br>**Viewshed Complexity: Density Quartile** | 21 | 1102.59 | 4.91 |
| **Season + Viewshed Complexity + Density Quartile + Season: Density Quartile** | 18 | 1110.63 | 12.95 |
| **Season + Viewshed Complexity + Density Quartile + Season: Viewshed Complexity +**<br>**Season: Density Quartile** | 24 | 1111.3 | 13.62 |
| **Season + Viewshed Complexity + Density Quartile + Season: Density Quartile +**<br>**Viewshed Complexity: Density Quartile** | 24 | 1114.22 | 16.54 |
| **Season + Viewshed Complexity + Density Quartile + Season: Viewshed Complexity +**<br>**Season :Density Quartile + Viewshed Complexity: Density Quartile** | 30 | 1115.43 | 17.75 |
| **Season + Viewshed Complexity** | 6 | 1127.05 | 29.37 |
| **Season + Viewshed Complexity + Season: Viewshed Complexity** | 12 | 1127.07 | 29.39 |
| **Viewshed Complexity** | 3 | 1128.28 | 30.6 |
| **Season + Density Quartile** | 7 | 1221.69 | 124.01 |
| **Season + Density Quartile + Season: Density Quartile** | 16 | 1231.63 | 133.95 |
| **Density Quartile** | 4 | 1231.68 | 134 |
| **Season** | 4 | 1262.47 | 164.79 |
| **None (intercept-only)** | 1 | 1273.64 | 175.59 |

<sup>a</sup>AICc is corrected Akaike's Information Criterion.
<sup>b</sup> $\Delta$ AICc is the difference between the two models being compared.
<sup>c</sup>We used this model in the analysis.

Estimates from the best-supported model of incidental detection varied from 0.07 (far distance quartile in high viewshed complexity) to 0.75 (near distance quartile in low viewshed complexity; Table 10). Thus, decoys placed in low viewshed complexity and closer to a turbine (i.e., in the near density quartile) had the highest probability of detection (0.75), whereas a decoy placed farther away from a turbine and in high viewshed complexity had the lowest probability of being detected (0.07; Table 10). The model describes a consistent pattern of decreasing incidental detection as distance from the turbine increases within each viewshed complexity class, and decreasing detection within each density quartile as viewshed complexity increases.

**Table 10.**
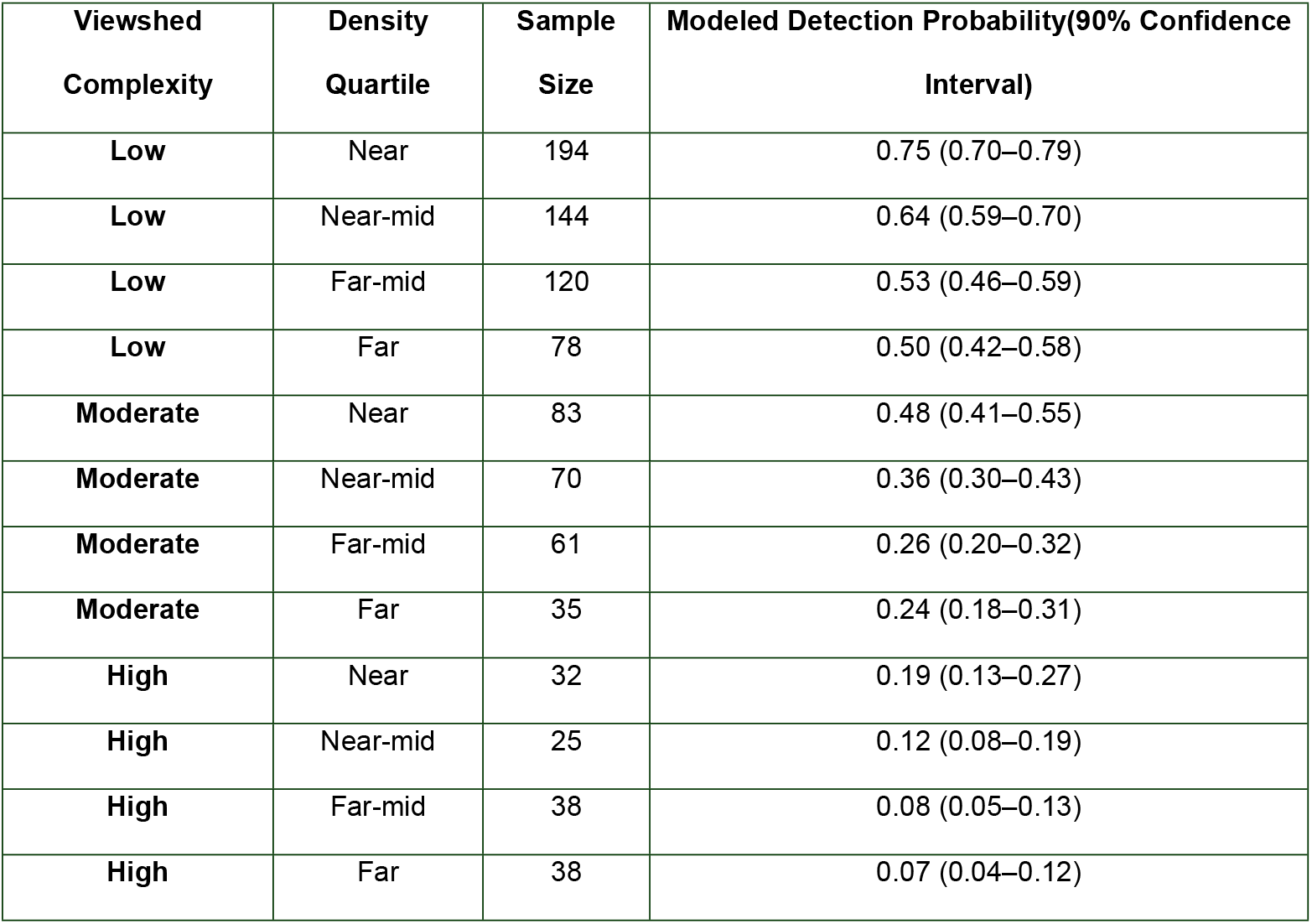
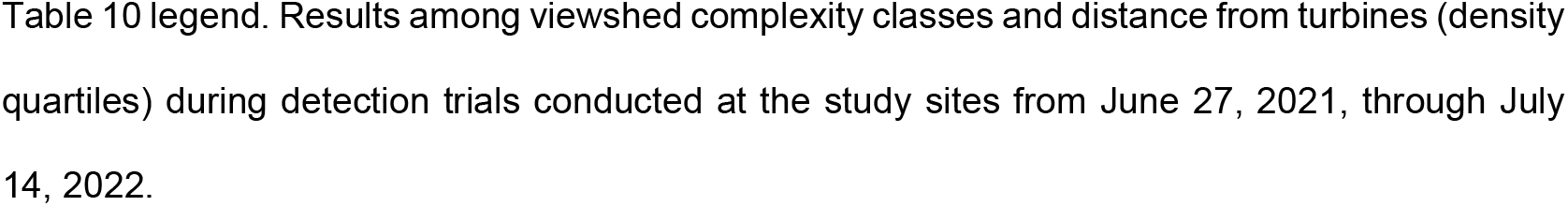
Incidental detection trial results among viewshed complexity classes and distance from turbines.

legend. Results among viewshed complexity classes and distance from turbines (density quartiles) during detection trials conducted at the study sites from June 27, 2021, through July 14, 2022.
| **Viewshed Complexity** | **Density Quartile** | **Sample Size** | **Modeled Detection Probability(90% Confidence Interval)** |
| --- | --- | --- | --- |
| **Low** | Near | 194 | 0.75 (0.70–0.79) |
| **Low** | Near-mid | 144 | 0.64 (0.59–0.70) |
| **Low** | Far-mid | 120 | 0.53 (0.46–0.59) |
| **Low** | Far | 78 | 0.50 (0.42–0.58) |
| **Moderate** | Near | 83 | 0.48 (0.41–0.55) |
| **Moderate** | Near-mid | 70 | 0.36 (0.30–0.43) |
| **Moderate** | Far-mid | 61 | 0.26 (0.20–0.32) |
| **Moderate** | Far | 35 | 0.24 (0.18–0.31) |
| **High** | Near | 32 | 0.19 (0.13–0.27) |
| **High** | Near-mid | 25 | 0.12 (0.08–0.19) |
| **High** | Far-mid | 38 | 0.08 (0.05–0.13) |
| **High** | Far | 38 | 0.07 (0.04–0.12) |

### Carcass persistence

The best-supported models of raptor persistence for each study site can be found in S2 Table. The best-supported models for Marble River and Wild Horse included a season covariate, while the models for all other sites were intercept-only. Median carcass persistence and average probability of persistence estimates (based on a 30-day search interval) for the datasets most representative of each of the study sites are presented below (Table 11). Persistence trial data from all 4 seasons were available to correspond with our incidental detection trials, with the exception of Wild Horse where summer data were not available. Instead, the fall estimate of persistence at Wild Horse was used for summer as a conservative proxy since it was the lowest of the 3 represented seasons. Median raptor persistence was typically in excess of the 30-day search interval assumed for O&M staff visitation to turbines, ranging from 19.46 days at the study site with mostly forest land cover (Marble River, informed by Arkwright Summit Wind Farm [Arkwright]) to 170.12 days at the study site with mostly cropland (Shiloh I and Frontier I, informed by Hale Wind Farm). The average probability of persistence, which is a component of *g*, was also high relative to a 30-day search interval, exceeding 0.85 in all but 1 case (0.61; Marble River, informed by Arkwright; Table 11). Model parameter estimates and 95% confidence intervals of the β parameter used in EoA to inform the incidental *g*’s can be found in S3 Table.

**Table 11.**
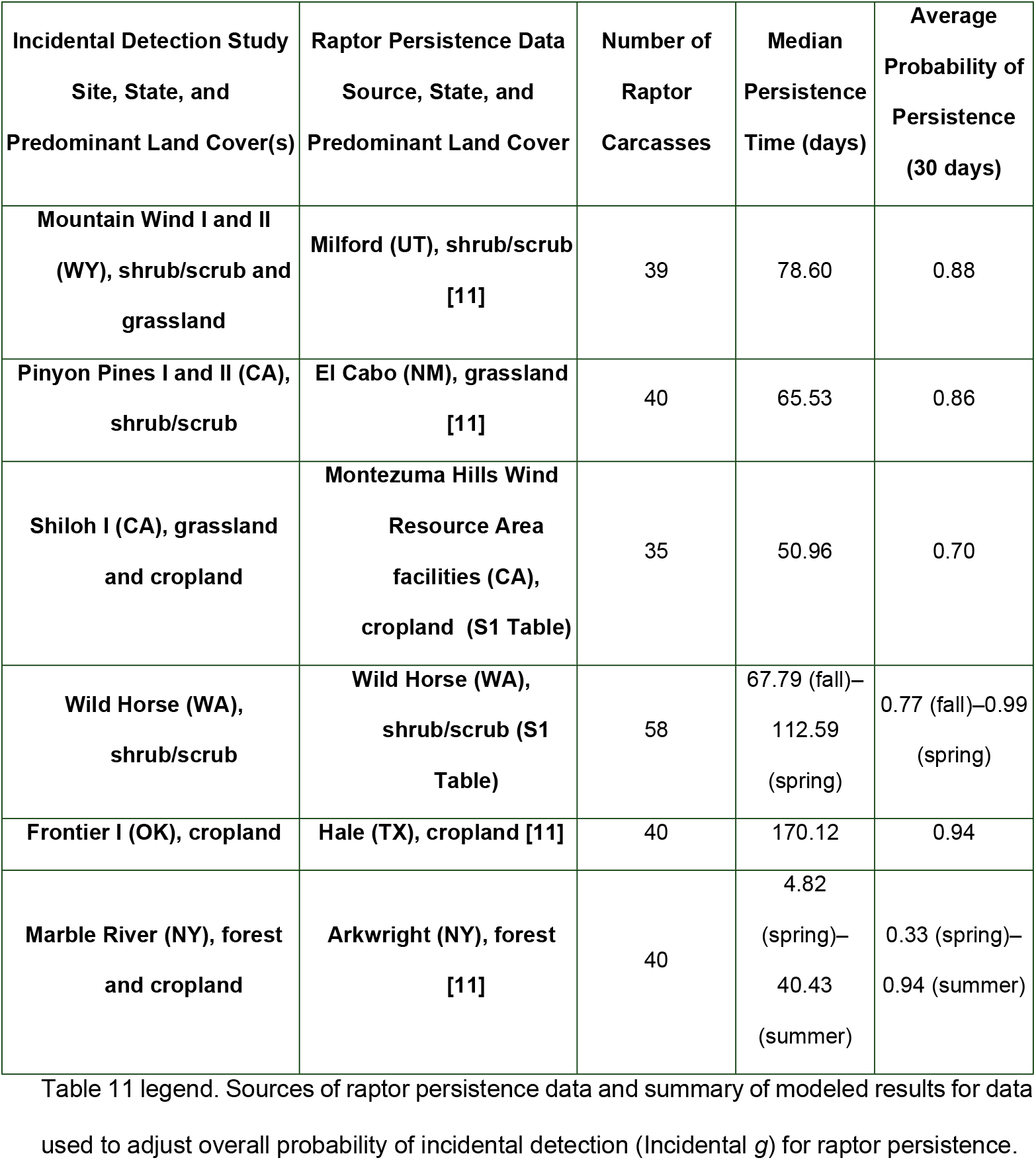
Sources of raptor persistence data and summary of modeled results.

### Area adjustment and proportion of turbines visited

The amount of viewable area at turbines searched (i.e., those we assumed could be visited by O&M staff) for each study site varied by season, with the exception of Pinyon Pines I and II and Mountain Wind I and II, where all areas within 100 m of a turbine were considered viewable in all seasons. Viewable area ranged from 0.18 (Marble River, summer) to 1.00 (Pinyon Pines I and II, all seasons studied; Table 12). Viewable area was lowest at study sites with cropland, particularly during the growing season (summer for our study sites). The proportion of viewable area during the non-growing season varied at each of the study sites, but not consistently. Marble River and Shiloh I increased viewable area by 18% and 13%, respectively, while Frontier I increased by 188%. Accounting for the predicted density of raptor fatalities relative to distance from turbine, the density-weighted area adjustment followed the same pattern as proportion of viewable area, ranging from 0.301 (Marble River, summer) to 0.939 (Pinyon Pines I and II, all seasons studied). In some cases, the density-weighted area adjustment was greater than the proportion of viewable area (e.g., Marble River, all seasons; Table 12) because viewable areas were generally at distances from the turbine base that aligned with the highest density of predicted raptor fatalities from the distribution model in Hallingstad et al. [10]. Finally, the proportion of turbines we assumed to have been searched by O&M staff during each season was generally 1.0 (i.e., all turbines visited), with seasonal exceptions for turbines located in areas dominated by agriculture (summer) or where access was precluded by snowfall (winter; Table 12). These subsets of turbines were considered to have effectively no searchable area while conditions prevented detection.

**Table 12.** Area adjustment for unviewable/unsearchable areas and proportion of viewable area.

are at turbines tested for incidental detection of eagles during detection trials conducted at the 6 study sites from June 27, 2021, through July 14, 2022.
| **Study Sites** | **Season** | **Predominant Viewshed** | **Proportion of Viewable**<br>**Area at Searched**<br>**Turbines<sup>a,b</sup>** | **Density-weighted Area**<br>**Adjustment for Searched**<br>**Turbines<sup>b,c</sup>** | **Proportion of Turbines**<br>**Searched** |
| --- | --- | --- | --- | --- | --- |
| **Frontier I** | Spring | Moderate | 0.857 | 0.662 | 1.00 (61/61) |
|  | Summer | Unviewable/unsearchable | 0.298 | 0.360 | 0.61 (37/61) |
|  | Fall | Low | 0.857 | 0.662 | 1.00 (61/61) |
|  | Winter | Low | 0.857 | 0.662 | 1.00 (61/61) |
| **Marble River** | Spring | Moderate | 0.216 | 0.342 | 1.00 (70/70) |
|  | Summer | Unviewable/unsearchable | 0.183 | 0.301 | 1.00 (70/70) |
|  | Fall | Moderate | 0.216 | 0.342 | 1.00 (70/70) |
|  | Winter | Unviewable/unsearchable | n/a | n/a | 0 (0/70) |
| **Mountain Wind I**<br>**and II** | Spring | Moderate | 0.978 | 0.932 | 1.00 (67/67) |
|  | Summer | Moderate | 0.978 | 0.932 | 1.00 (67/67) |
|  | Fall | Moderate | 0.978 | 0.932 | 1.00 (67/67) |
|  | Winter | Unviewable/unsearchable | n/a | n/a | 0 (0/67) |
| **Pinyon Pines I**<br>**and II** | Spring | High | 0.999 | 0.939 | 1.00 (100/100) |
|  | Summer | High | 0.999 | 0.939 | 1.00 (100/100) |
|  | Fall | High | 0.999 | 0.939 | 1.00 (100/100) |
|  | Winter | High | 0.999 | 0.939 | 1.00 (100/100) |
| **Shiloh I** | Spring | Moderate | 0.732 | 0.775 | 1.00 (100/100) |
|  | Summer | Moderate | 0.677 | 0.752 | 0.59 (59/100) |
|  | Fall | Low | 0.732 | 0.775 | 1.00 (100/100) |
|  | Winter | Moderate | 0.732 | 0.775 | 1.00 (100/100) |
| **Wild Horse** | Spring | Moderate | 0.899 | 0.873 | 1.00 (149/149) |
|  | Summer | Moderate | 0.899 | 0.873 | 1.00 (149/149) |
|  | Fall | Moderate | 0.899 | 0.873 | 1.00 (149/149) |
|  | Winter | Moderate | 0.899 | 0.873 | 1.00 (149/149) |

<sup>a</sup>Cropland areas were considered unviewable/unsearchable during the summer season at Frontier I, Marble River, and Shiloh I; detection in these areas was assumed to be 0 until harvest occurred.
<sup>b</sup>n/a = not applicable; during periods where Operations and Maintenance staff were not routinely traveling the study site, detection trials were paused and it was assumed no areas were searched until routine travel resumed.
<sup>c</sup>The density-weighted area adjustment can exceed the viewable proportion of the search area when carcass distribution is expected to be relatively high within viewable areas (see Hallingstad et al. [10]).

### Overall probability of incidental detection (Incidental *g*)

The resulting incidental *g* for the 12-month period for our study sites ranged from 0.07 (Marble River) to 0.47 (Mountain Wind I and II; Table 13). There was variability in incidental *g* by season at each study site, with the greatest range occurring at Frontier I (0.12–0.50), and the smallest range occurring at Marble River (0.06–0.11, excluding winter; Table 13). Given the long persistence times for raptors based on the persistence models used at 5 of our 6 study sites, the primary drivers of variability in incidental *g* were the detection probability and area adjustment. At Frontier I, the incidental *g* was lower during the active crop period (0.12 in summer) than other seasons (0.25 to 0.50; Table 13), but the effect was decreased by the insensitivity of the density-weighted area adjustment to reduced viewable area. For both Shiloh I and Frontier I, higher viewshed complexity during the spring season resulted in lower detection probabilities and, therefore, decreased overall probabilities of detection in this season (0.24 and 0.25, respectively) relative to fall and winter. Pinyon Pines I and II had its lowest incidental *g* in winter, when detection probability was lowest, while Wild Horse had its highest incidental *g* in spring, when both carcass persistence and detection probabilities were highest. At Marble River, the incidental *g* was lowest during spring due to reduced raptor carcass persistence during that season (see Table 11). Mountain Wind I and II had relatively consistent incidental *g*’s between the 3 seasons in which detection trials were conducted (Table 13), as this study site had relatively consistent detection probabilities, persistence estimates, and searchable area in spring, summer, and fall.

**Table 13.**
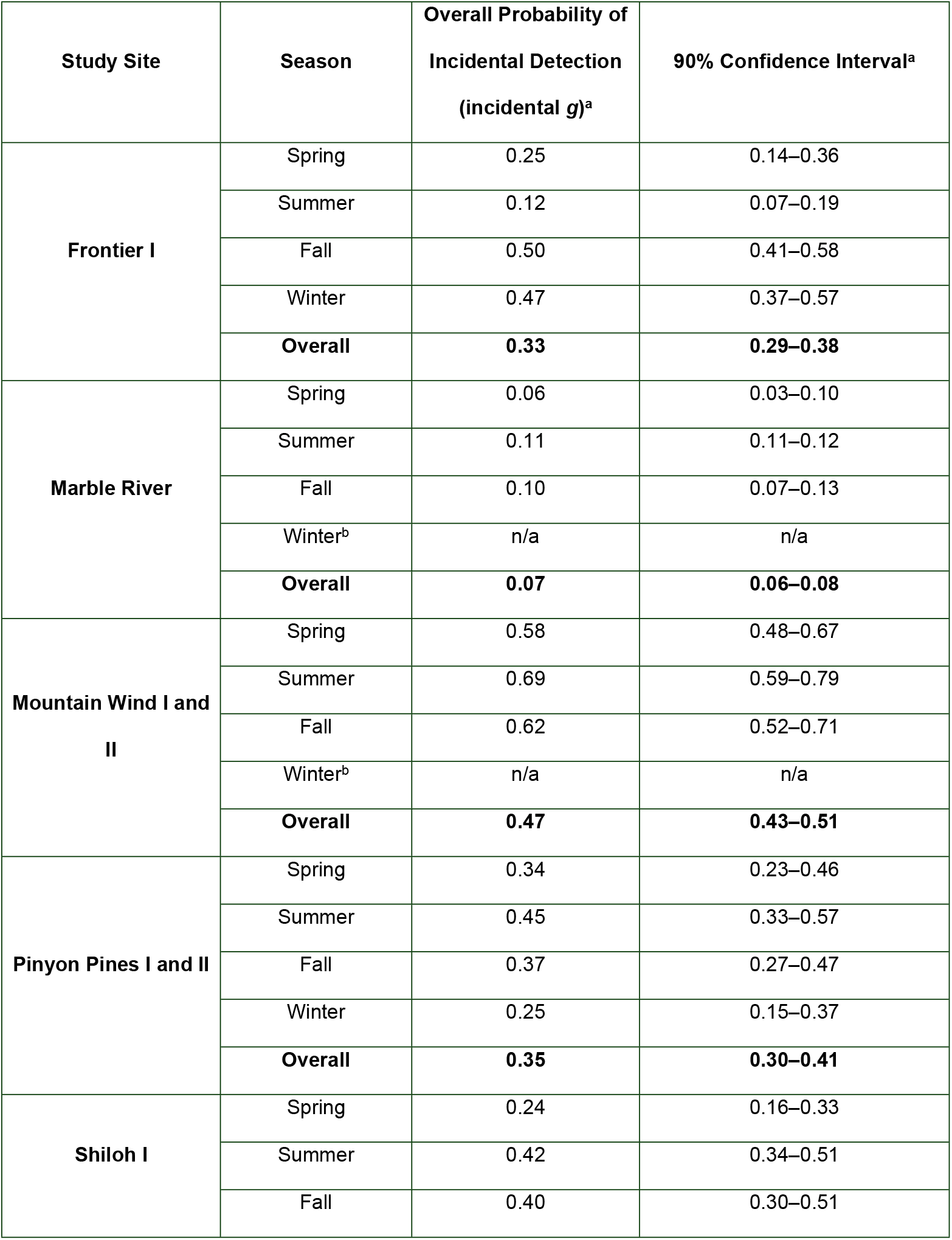
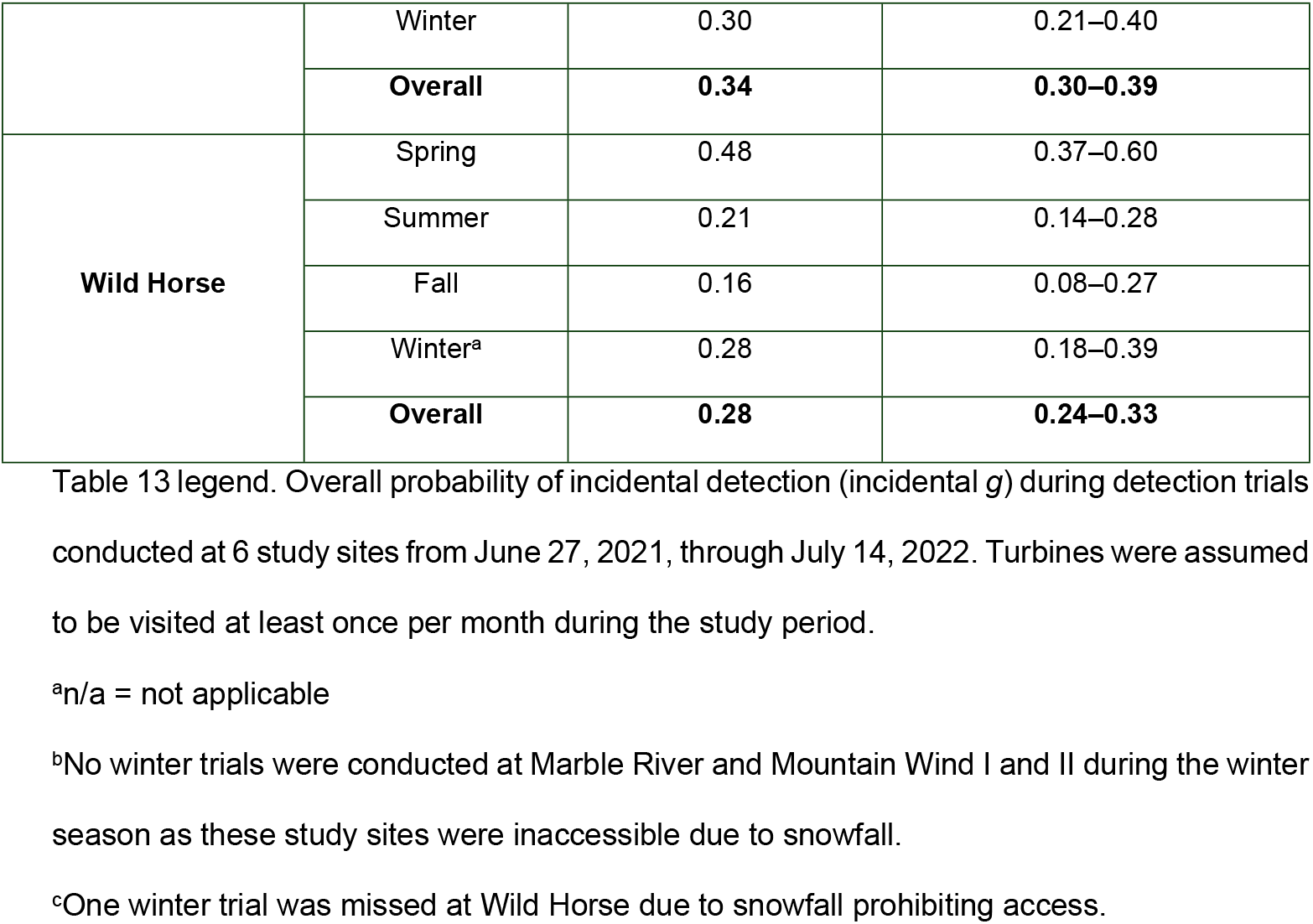
Overall probability of incidental detection.

## Discussion

We found O&M staff are successful at detecting large carcasses. O&M staff making regular turbine visits were capable of incidentally detecting approximately 1 out of every 2 decoys at our study sites, on average. The probability of O&M staff detecting decoys exceeded 0.25 at all study sites, reaching as high as 0.78. In a review of publicly available data from wind energy facilities, Bay et al. [35] found searcher efficiency ranged from roughly 0.43 to 1.0 for large bird carcasses (e.g., mallards, rock pigeons [*Columba livia*]) in 26 fatality monitoring studies administered by third parties, with a mean searcher efficiency of approximately 0.74. Our results indicate incidental detection, under ideal conditions, can approach detection probabilities resulting from eagle fatality monitoring studies employing transect and other more time-intensive search methods. Further, our results are congruent with a review of eagle fatalities in which the majority of eagle carcasses were detected incidentally during routine activities at wind energy facilities [3].

The true benefit of incidental detection requires calculation of *g*, which facility operators can then use to calculate fatality estimates and evaluate permit compliance. The *g* accounts for other sources of bias inherent with fatality monitoring, including the probability of a carcass persisting through the search interval and the proportion of carcasses that occur within searched areas. The annual incidental *g* at each study site ranged from 0.07 to 0.47, with seasonal detection probabilities ranging from 0 to 0.69. Our study sites covered a wide geographic range and included a variety of landscape conditions representative of many wind facilities. Combining all study sites, the average incidental *g* in our study was 0.31. In some circumstances, incidental *g* approached or exceeded *g*’s attainable through traditional standardized search methodology used in general bird and bat fatality monitoring (e.g., 30% of turbines searched out to tip-height radii using 6-m transect spacing results in a maximum *g* of 0.30; see Strickland et al. [36]).

Our study shows that quantifying the incidental *g* is possible and can contribute to a better understanding of a facility’s direct impacts on eagles. A potential advantage of incidental detection is reduced third-party monitoring burden placed on facility operators. Standardized searches are labor intensive, particularly if prescriptive monitoring requirements for regular searches of large areas underneath all turbines were necessary to meet the *g* target for compliance under an ETP [16-19]. Traditional monitoring often requires at least 2 hours of labor per turbine search and mobilization of a third party for monitoring [36]. Another advantage of incidental detection is that O&M staff are on site for the life of the facility, potentially offering long-term information on eagle and other large raptor mortality at all operating wind facilities. An improved understanding of eagle mortality would assist wildlife agencies and researchers aiming to evaluate how installed and future wind energy build-out could affect population trends of raptors [37-39].

### Evaluating influences on incidental detection

Viewshed complexity was 1 of 2 primary drivers of the ability of O&M staff to detect decoys placed in our study. Not surprisingly, incidental detection probability decreased as viewshed complexity increased. Decoys placed in low viewshed complexity sites were detected twice as often as those placed in moderate viewshed complexity sites, and detected 6 times more often than those placed in high viewshed complexity sites (Table 9). Even when considering distance from turbine, detection probabilities for decoys were consistently higher in low viewshed complexity sites than in moderate viewshed complexity sites. In other words, a decoy placed more than 61 m from a turbine base in low viewshed complexity had a greater detection probability than a decoy placed less than 33 m from a turbine base in a moderate viewshed complexity. Similarly, detection probability within moderate viewshed complexity was higher within every density quartile than detection in any high viewshed complexity density quartile (Table 10). Incidental detection probability should therefore be highest at facilities with predominately low viewshed complexity within 100 m of turbine bases. Although season was not a covariate in the most parsimonious detection probability model, we acknowledge viewshed complexity and season are correlated. Our analysis shows that viewshed complexity better explains variation in incidental detection across the entire dataset, an intuitive result as detection probability would not be driven by calendar date. Nonetheless, seasonal changes in viewshed complexity and area adjustment (see below) are important to consider when determining potential influences on incidental detection.

Distance from turbine (as represented by the density quartile variable) was the second main factor explaining detection probability by O&M staff. Again, not surprisingly, decoys placed farther from turbine bases were less likely to be detected, a trend consistent with the distance effect on detection we found when testing a scanning search methodology [10]. This pattern was consistent within each viewshed complexity class; however, detection probabilities across viewshed complexities overlapped by density quartile. For example, detection probabilities within the far density quartile (more than 61 m from turbine bases) ranged from 0.07 to 0.50, whereas detection within the near density quartile (less than 33 m from turbine bases) ranged from 0.19 to 0.75 (Table 10). These results suggest viewshed complexity has a larger impact on incidental detection than distance from turbine.

When O&M staff reported the activity they were participating in when a detection occurred, almost a third of all decoy detections were made while driving through the study sites. Eagle carcasses are large and can be highly visible on the landscape in certain conditions; it is important that area adjustment calculations include all areas within a minimum of 100-m of turbine bases that are visible from roads traveled by O&M staff. As turbine sizes increase and the distribution of carcasses may become more widely spread under larger turbines, operators of new facilities may want to extend the search area along roadsides to account for detections made in these areas by O&M staff while they are driving. Another 30% of detections occurred during unscheduled maintenance activities at turbines, either while technicians were on the turbine pad or while they were “up tower.” Although we did not assess viewsheds from tower nacelles, it is possible that some proportion of these maintenance activity detections occurred from the nacelles. Encouraging the practice of a quick scan of the ground within 100 m of turbine bases while technicians are up-tower may be another way to increase the incidental *g*.

A critical component of effective incidental monitoring is to ensure O&M staff are educated and adequately trained in fatality detection and reporting procedures. Operators should encourage heightened awareness of the benefits for incidental detection as an effective tool at their facilities to ensure compliance with ETP conditions while reducing the need for third-party involvement. Encouraging O&M staff awareness and diligence through proper training (e.g., developing a “search image”) and incentivization can foster a culture of responsibility and O&M staff satisfaction in their ability to contribute towards ETP compliance. We found that O&M data were not reported consistently at each site throughout the entire year, limiting our ability to incorporate some data into our modeling efforts (e.g., O&M activity when detection occurred). Data reliability can be enhanced through periodic training and by testing carcass detection by O&M staff with detection trials administered by a third party.

### Other influences on the overall probability of incidental detection (incidental *g*)

Incidental *g* can be affected by variability in incidental detection, as well as patterns in carcass persistence and the area adjustment. Areas not visible from turbine bases or surrounding roads reduce incidental *g* by requiring an area adjustment; the magnitude of this effect will depend upon how far the unviewable/unsearchable areas are from the turbine bases. For example, if topography blocks view of an area more than 61 m from a turbine base, the impact on detection will be mitigated by reduced large carcass density at such distances (i.e., the unviewable/unsearchable area in this example would be expected to contain fewer carcasses than areas closer to the turbine base).

Seasonal effects can also influence detection probability by decreasing the effective search area. Snowfall and continual snow cover precluded O&M staff from adhering to their regular turbine visitation schedule at 3 of our study sites. When O&M staff are not routinely traveling the facility because conditions are not conducive to travel, detection is expected to be 0. Carcasses can be found once the snow melts, but quantifying the effect of such conditions on carcass condition and detectability fell outside the scope of this study.

Another seasonal effect was seen when crops reach heights that prevented detection, which we treated as unsearchable/unviewable areas in our study. Wind facilities in predominately agricultural areas will face a similar challenge during the peak growing season as facilities dealing with winter closures. The impact on detection will be dictated by the length of these periods, and in the case of cropland, the extent of the cropland areas and the density of carcasses estimated to occur within them (as based on carcass distribution models). For some facilities, eagle risk may not overlap these challenging seasonal conditions, so facilities evaluating the potential of incidental detection should focus on ground cover conditions during the seasons where collision risk is possible.

Carcass persistence is also an important factor in *g*. In our study, we used the best available raptor persistence data. For 5 of our 6 study sites, this required applying persistence data from surrogate facilities in the same geographic areas and with similar land cover types. We assume the persistence data used in our analyses are representative of what we would observe if raptor persistence trials were conducted at these study sites. In the case of Marble River, we used persistence data from another northeastern wind facility located in forest habitat. The resulting probability of a carcass persisting through a 30-day interval at Marble River was 0.33 during the spring season; this resulted in the incidental *g* during spring being 40% lower than during summer at Marble River despite having higher incidental detection (0.77 versus 0.57) in spring and a lower proportion of viewable area (0.301 versus 0.342) in the summer. Decreasing the search interval would result in more carcasses persisting until the next search round, and would have a pronounced effect at facilities with low carcass persistence. If practicable, an operator could increase the incidental *g* at their facility by implementing more frequent turbine visitation (e.g., twice monthly).

### Using the overall probability of incidental detection (incidental *g*) in fatality estimation

Our study shows incidental detection can be an effective and efficient tool for gathering fatality monitoring data necessary to quantify eagle fatality rates and help permittees document compliance with permitted take. Using EoA, researchers can use the incidental *g* to adjust the number of eagles found to an estimated number of eagle fatalities during years in which standardized fatality surveys are not conducted, or to calculate an average *g* for a multi-year period during which standardized monitoring occurs in some years but others. If we apply the EoA estimator for a facility where O&M staff conducting monthly turbine visits incidentally found 1 eagle carcass over a 5-year permit term, and we use estimated incidental *g*s of 0.10, 0.20, 0.30, 0.40 and 0.50, this would result in 12, 6, 4, 3, and 2 estimated eagle fatalities, respectively (Table 14). If 2 carcasses are found during a year, this would result in 22, 11, 7, 5, and 4 estimated eagle fatalities, respectively (Table 14). A permittee’s ability to utilize the contribution of incidental detection to estimate eagle fatality rates in years without standardized eagle fatality monitoring has practical implications with respect to demonstrating eagle take permit compliance, quantifying the level of mitigation required, and likelihood for adaptive management actions if the facility were to exceed permitted take. Failing to factor in the contributions of incidental detection could lead to artificially inflated take estimates, potentially resulting in a permittee implementing costly and unnecessary measures to reduce take, additional mitigation requirements, and even permit suspension or revocation if permit take limits are exceeded.

**Table 14.** Estimated eagle fatalities based on overall probability of incidental detection.

| Overall Probability<br>of Detection (*g*) | Eagle<br>Carcasses<br>Found = 0 | Eagle<br>Carcasses<br>Found = 1 | Eagle<br>Carcasses<br>Found = 2 | Eagle<br>Carcasses<br>Found = 3 | Eagle<br>Carcasses<br>Found = 4 |
| --- | --- | --- | --- | --- | --- |
| **0.10** | 2 | 12 | 22 | 32 | 42 |
| **0.20** | 1 | 6 | 11 | 16 | 21 |
| **0.30** | 0 | 4 | 7 | 10 | 14 |
| **0.40** | 0 | 3 | 5 | 8 | 10 |
| **0.50** | 0 | 2 | 4 | 6 | 8 |

Facility operators must evaluate ETP requirements when determining the value of incidental detection for their circumstances and determine the appropriate use of incidental detection versus standardized fatality monitoring efforts used for fatality estimation. Our study shows incidental detection is a valuable tool that can supplement or, in some conditions, fully replace standardized fatality monitoring efforts.

### Evaluating the potential for incidental detection at a wind facility

We believe our results show incidental detection can be part of a viable eagle fatality monitoring study design for ETPs, but facility operators should carefully evaluate their facility to assess the potential for incidental *g* to meet their monitoring objectives. This assessment involves a few simple steps. First, we recommend viewshed mapping as a preliminary step in assessing the viability of incidental detection at a given facility. Areas within 100 m of turbine bases that are not visible from pads or roads, or have crop height and density that preclude detection (i.e., unviewable/unsearchable), will reduce the effective search area. Areas with high viewshed complexity, such as a tall, thick scrub-shrub plant community, will also reduce detection probability. The effect of lower detection probability on *g* will be larger if carcass density is expected to be high (i.e., areas within 61 m of turbine bases) within difficult to search or unsearchable areas. Conversely, if high viewshed complexity areas are more than 61 m from turbine bases, the effect of lower detection probability on *g* will not be as strong because less than 25% of carcasses are expected within these areas. We suggest that operators also consider winter access limitations at their facilities, as incidental detection can only occur when O&M staff are traveling to or visiting turbines. Seasonal viewshed mapping may be needed if substantial access and vegetation changes occur throughout the year.

Next, operators aiming to incorporate incidental detection probability into fatality estimation should be sure to maintain (or implement) a regular turbine visitation schedule. For our study, each study site had a standard operating procedure involving at least 1 visit to every turbine each month. This is a typical maintenance check schedule for wind energy facilities, but is not applied universally. EoA requires an assumed search interval, as this ties into the probability of a carcass persisting until the next opportunity for detection. If practicable, an operator may also choose to increase turbine visitation (i.e., decrease their search interval) if the eagle take permit threshold supports maximizing detection. We also encourage all operators to use the best available raptor carcass persistence data to determine the appropriate carcass persistence estimate for their facility; in the absence of suitable raptor carcass persistence data, site-specific game bird persistence data can be adjusted and used in eagle fatality estimation [11].

### Implications and next steps

Our study resulted in the successful application of a general field methodology in which an eagle carcass surrogate distributed throughout a facility can be used to measure the incidental *g* resulting from O&M staff performing their regular activities. Furthermore, the resulting incidental *g*’s support the inclusion of incidental detection into fatality monitoring study designs and fatality estimation. Implementing a combination of standardized and incidental monitoring with the appropriate data quality standards, particularly in non-standardized monitoring years, can provide an efficient and viable fatality monitoring approach to assess consistency with ETP terms and conditions. Furthermore, at facilities with conditions that support high detection probability (e.g., flat topography, low/sparse vegetation), standardized monitoring may not be required to meet compliance with ETP conditions.

Opportunities exist to refine our understanding of incidental carcass detection. First, we assumed decoys are a reasonable surrogate to evaluate detection of eagle carcasses. Eagle carcasses and parts are protected by federal law and are prioritized for Native American religious purposes, so obtaining authorization to use eagle carcasses for bias trials is unlikely. In the absence of eagle carcasses, researchers may choose to evaluate the suitability of decoys as a surrogate for testing detection by comparing detection probabilities between decoys and large raptor carcasses. Second, EoA incorporates a detection reduction factor (*k*) that describes how carcass detection changes between searches. In our study, we assumed a probability of 0.67 for this factor, which is the only published *k* currently available [29]. We did not evaluate the detection of partial carcasses and feather spots; we acknowledge this as a potential source of bias for these trials, because partial carcasses and feather spots likely have different detection probability compared to intact decoys. However, feather spots can often cover a much larger area than a decoy or an eagle carcass, resulting in an increased detection probability at this stage of scavenging and decomposition. Furthermore, of the 103 eagle fatalities in WEST’s Renew database for which “physical condition” of the fatality was reported, 81 carcasses (79%) were at least partially intact and only 7 (7%) were found as feather spots [40]. Yet, we do not know the factor by which eagle detection changes between searches, and suggest further studies to estimate these factors for the purpose of refining fatality estimates. Lastly, we measured incidental detection under a variety of landscape contexts and environmental conditions throughout a single year. However, incidental detection has been little studied and we documented a wide range of overall probabilities of detection in this study. Additional research will strengthen our inferences and add to our understanding of factors influencing incidental detection probability.

## Acknowledgements

Will Vesely and Juan Botero served as the Renewable Energy Wildlife Research Fund’s (REWRF) project managers and provided valuable administrative support during our research. We would like to thank the O&M staff members at our 6 study sites for participating in the study, and for providing safety instruction, access, storage, and other logistical support. H.T. Harvey & Associates and NextEra provided valuable raptor persistence data for the Shiloh I area. Andrea Palochak and David Klein (both with WEST) provided valuable assistance during manuscript preparation. Faith Kulzer (with WEST) provided support organizing, formatting, and preparing data for analyses. We thank Paul Rabie, Amanda Hale, and Karl Kosciuch (all with WEST), who offered thoughtful comments on our initial manuscript submittal.

## Supporting information captions

**S1 Table. Large raptor carcass persistence dataset**. Table includes raw data necessary to replicate carcass persistence analyses, from persistence trials not publically available or previously published. Data for Shiloh I was collected from nearby wind facilities in the Montezuma Hills Wind Resource Area. Data for Wild Horse was collected in previous years at Wild Horse.

**S2 Table. The best-supported survival regression models of large raptor carcass persistence for each study site**.

**S3 Table. Model distribution, and required parameter estimates and 95% confidence intervals of the survival regression parameter(s) used in Evidence of Absence to inform the overall probability of incidental detection (incidental *g*) estimates**.

